# Engineering non-exponential proliferation in *Escherichia coli* using functionalized protein aggregates

**DOI:** 10.1101/2025.08.23.671898

**Authors:** Ronald Van Eyken, Sam Jordens, Dries Oome, Kevin Broux, Abram Aertsen

## Abstract

Uncurbed exponential proliferation might not always be required for genetically modified microorganisms, and might even cause unpredictable liabilities in their behavior and impact. We therefore constructed a bacterial chassis that adheres to linear proliferation for a finite number of generations, based on the asymmetric segregation of a gradually declining intracellular protein aggregate that is engineered to produce a conditionally essential metabolite such as cAMP.

## INTRODUCTION, RESULTS and DISCUSSION

Exponential proliferation is innate to microbial life, and microbial populations can rapidly expand under permissive conditions [1]–[3]. However, with the increasing potential and envisaged applications of genetically modified microorganisms (GMM), the brute force of exponential growth might not always be required and might even cause unpredictable liabilities in the behavior and impact of GMMs. Indeed, prolific expansion of e.g. therapeutic GMMs *in vivo* could challenge biocontainment efforts and make the intended dose and duration of the therapy highly unpredictable. As such, deterministically engineering a microbial chassis away from exponential growth, and thus fundamentally reprogramming population growth dynamics, is a relevant challenge in synthetic biology.

In this context, an interesting example of curbed microbial growth in nature occurs in *Mycobacteria smegmatis*, where the cell’s elongation complex is exclusively active at the old cell pole [4]. Asymmetric segregation of this elongation complex among daughter cells upon cell division gives rise to faster growing accelerator cells (inheriting the existing elongation complex) and slower growing alternator cells (that first need to rebuild and activate their own elongation complex) [5]. Furthering on this natural concept, linear (as opposed to exponential) proliferation could be obtained if the cell’s ability to grow would depend on a feature that strictly segregates asymmetrically between daughter cells, creating a growing (founder) cell and a non-growing cell. Moreover, if this feature itself would autonomously diminish and disappear over time, a founder cell could only perform a limited number of divisions.

Over the past years, a number of proteins (e.g. Tsr as part of the chemotaxis complex [6]) and protein assemblies (such as protein aggregates and protein condensates [7]–[9]) have been shown to segregate asymmetrically. We therefore decided to engineer an intracellular protein aggregate (PA) with the metabolic capacity to produce cAMP, as a key signaling molecule for *Escherichia coli* to be able to grow on secondary carbon sources [10], [11]. More specifically, a plasmid (i.e. pAJM657-2PA) was constructed from which two tri-partite fusion proteins (i.e. T18-mCer3-cI78^EP8^ and T25-mSc-I-cI78^EP8^; Fig. 1A) could be conditionally expressed. Each of these fusion proteins contains (***i***) an inert part of the catalytic domain of the *B. pertussis* CyaA protein (i.e. T18 or T25) that only in close vicinity with the other part reconstitutes cAMP synthase activity [12], [13], (***ii***) a fluorescent protein (i.e. mCerulean3 or mScarlet-I [14], [15]) that allows to microscopically track and validate the intracellular whereabouts of the fusion proteins, and (***iii***) a previously characterized autonomously aggregating cI78^EP8^ moiety [16] that forces the fusion proteins to co-aggregate (Figs. 1A and 1B). Importantly, exclusive reconstitution of the split cAMP synthase on the PA (i.e. via the cI78^EP8^-mediated co-aggregation of T18 and T25) ensures that cAMP emerges only from the PA and not from proteins or protein parts that are released from the PA by chaperones or proteases. This pAJM657-2PA plasmid was subsequently transformed to *E. coli* MG1655 *ΔcyaA ΔclpB lacY-syfp2* that lacks its own cAMP synthase (*ΔcyaA*), lacks the ClpB disaggregase in order to stabilize PAs (*ΔclpB*) [17]– [19], and carries the *lacY-syfp2* transcriptional reporter so that cAMP-dependent expression of its *lacZYA* locus can be fluorescently monitored [20].

**Figure 1:**
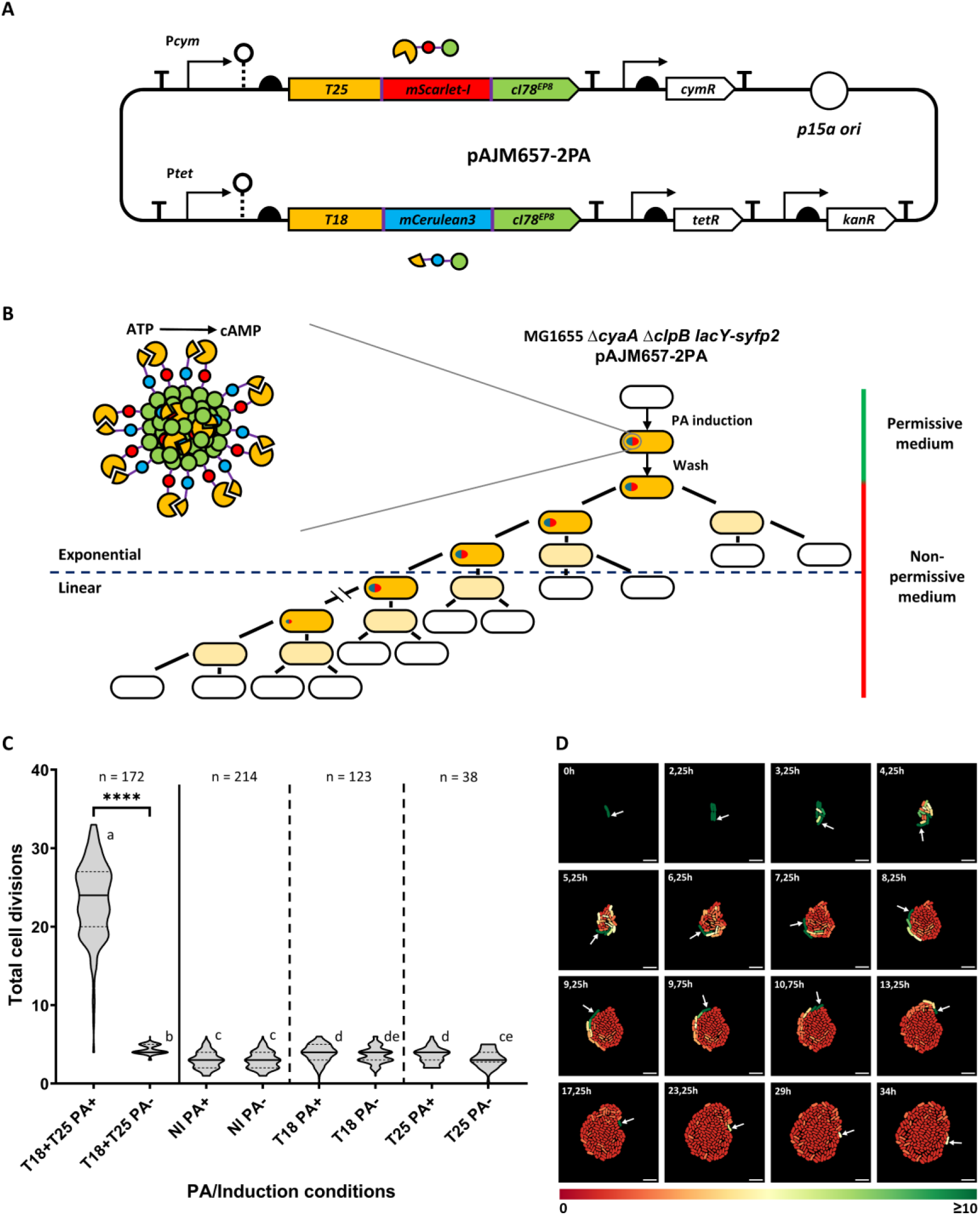
(A) Functional map of the pAJM657-2PA plasmid encoding the T18-mCer3-cI78^EP8^ and T25-mSc-I-cI78^EP8^ fusion proteins. (B) Schematic representation of the envisioned synthetically curbed propagation of *E. coli*, based on the asymmetric segregation of a cAMP generating PA. (C) Counted total cell divisions for individual MG1655 *ΔcyaA ΔclpB lacY-syfp2* pAJM657-2PA cells in a 48h TLFM experiment on non-permissive (ABc) medium per PA/induction condition displayed as truncated violin plots (cured datasets). PA+ indicates daughter cells inheriting the PA bearing cell pole, whereas PA- indicates daughter cells inheriting the PA free cell pole from the first PA+ mother cell. T18-mCer3-cI78^EP8^ and T25-mSc-I-cI78^EP8^ fusion proteins were either both (T18+T25) or separately (T18/T25) induced or not induced (NI). The number of tracked cells is indicated by n. Violin plots with non-corresponding letters are significantly different from one another (p-value ≤ 0.05, determined by Conover-Iman test with Bonferroni correction after rejection of the null hypothesis of a Kruskal-Wallis test). Four asterisks indicate a p-value ≤ 0.0001. (D) Consecutive time-lapse images of a representative microcolony of the T18+T25 condition, false colored by in-house image analysis to visualize future growth potential of each cell in the microcolony. Color represents amount of progeny in the next 15h of growth. White arrows indicate PA inheriting daughter cell. Scale bar corresponds to 5 µm.

Under non-induced control conditions, or when inducing only the T18-mCer3-cI78^EP^ or the T25-mSc-I-cI78^EP8^ fusion protein, the cells did not express *lacY-syfp2* (indicating their inability to synthesize cAMP) and were correspondingly growth deficient on amino acids as carbon-source (Fig. 1C, Extended Data Fig. 3, supplemental Movies S2, S3 and S4). However, transient pulse-expression of both T18-mCer3-cI78^EP8^ and T25-mSc-I-cI78^EP8^ led to cI78^EP8^-mediated co-aggregation of both fusion proteins into a single mCerulean3 and mScarlet-I labelled PA (Extended Data Fig. 3). Moreover, on this resulting PA, the T18 and T25 moieties became positioned in close enough proximity to enable cAMP synthesis (as inferred by *lacY-syfp2* activation in PA-bearing cells) and corresponding growth on amino acids as carbon source (Fig. 1C, Extended Data Fig. 3). Subsequent asymmetric segregation of this PA then allowed the PA-bearing (and thus cAMP producing) sibling to keep growing on amino acids, while continuously spawning off PA-lacking siblings that soon abrogated growth as their cAMP levels and related proteins cytoplasmically depleted (Fig. 1D, supplemental Movie S1). In fact, for the tracked PA-bearing cells, an average of ca. 23 divisions could be observed (Fig. 1C), which is likely still an underestimation because of progressive depletion of the agarose medium. In turn, cells not inheriting the PA stopped growing after ca. 4 divisions on average, which is similar to non-induced control cells (Fig. 1C).

As expected, the size of the initial PA was positively correlated with the number of divisions it endowed to the PA-bearing cell (Extended Data Fig. 2B, C). Moreover, since PAs inevitably become disaggregated over time, PA bearing cells eventually lose their PA and thus their ability to produce cAMP and proliferate on amino acids as carbon source. In fact, restoring the *clpB* gene within the host (creating *E. coli* MG1655 *ΔcyaA lacY-syfp2*) expedited PA disaggregation, and therefore clearly downtuned PA-sizes together with the average number of divisions of PA-bearing cells (Extended Data Fig. 5A, B). Synthetically setting the expression level of *clpB* can therefore serve as a tuner for the average number of divisions of the chassis.

Based on the above engineered principle of an asymmetrically segregating (and gradually decaying) enzyme producing a symmetrically diluting essential metabolite, an individual-based model of the constructed chassis confirmed linear growth dynamics in which only a fixed amount of (PA-bearing) founder cells consistently partake in growth (Fig. 2A, B). Moreover, the extrapolation of this model towards bulk population growth could in turn be experimentally confirmed. In fact, the growth dynamics of a population of PA-bearing founders as monitored via bulk optical density measurements indeed revealed the predicted linear growth profile (Fig. 2C, Extended Data Fig. 4).

**Figure 2:**
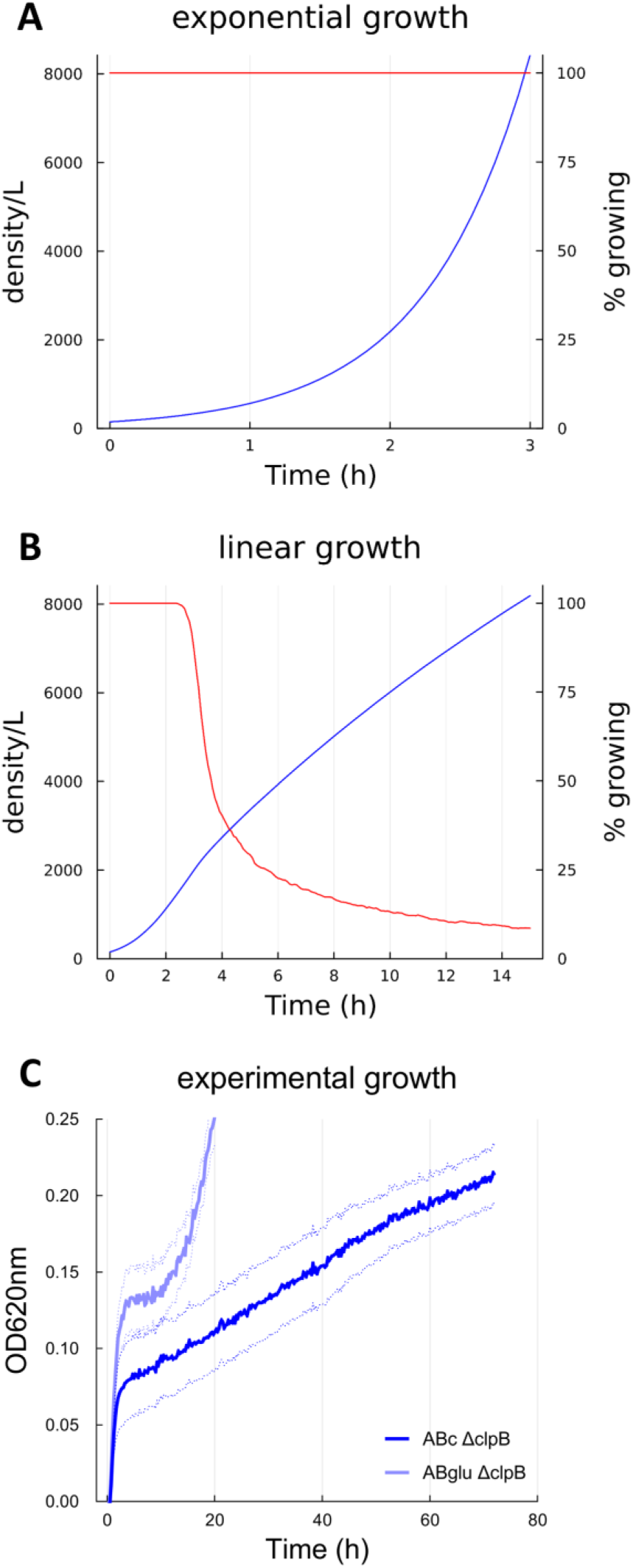
(A and B) Predicted growth dynamics for natural exponential growth (over 3h, A) and engineered linear growth (over 15h, B). Blue lines indicate total cell count (left axis), while red lines indicate the percentage of cells within the population that is still able to grow (right axis). (C) Optical density (OD_620_) monitored bulk growth profile of ca. 52000-96000 initial cells of MG1655 *ΔcyaA ΔclpB lacY-syfp2* pAJM657-2PA (with prior pulse induction of both T18-mCer3-cI78^EP8^ and T25-mSc-I-cI78^EP8^ fusion proteins) in either permissive ABglu (light blue line showing exponential growth) or non-permissive ABc (dark blue line showing linear growth) medium, together with the corresponding standard deviation (dotted lines).

In summary, we have demonstrated that protein aggregation tags (such as the cI78^EP8^ moiety) can be used to co-aggregatively display proteins in close enough vicinity to reconstitute a split enzyme that subsequently produces a conditionally essential metabolite. Since the resulting and asymmetrically segregating PA is the only source of this metabolite, only the founder cells inheriting this PA can keep growing and dividing. In turn, this results in a linearly proliferating population that halts growth when this PA becomes fully disaggregated. This proof-of-concept chassis can spur further developments of GMMs whose proper action and/or biocontainment could become compromised by uncurbed exponential growth.

## MATERIALS AND METHODS

### Strains and growth conditions

All bacterial strains and plasmids that were employed in this study are listed in Extended Data Table 1. Throughout the study, *Escherichia coli* (K12) MG1655 was used as a background for strain engineering. Bacteria were cultured either in LB (5 g/L NaCl [Fisher chemical], 5 g/L yeast extract [Oxoid], 10 g/L casein peptone type I [Neogen], and - in case of solid medium - 15 g/L of agar bacteriological No. 1 [Neogen]) or AB medium (ABglu; (1/5 diluted 5xA solution (10 g/L (NH_4_)_2_SO_4_ [Acros organics], 37.5 g/L Na_2_HPO_4_(H_2_O)_2_ [VWR BDH chemicals], 15 g/L KH_2_PO_4_ [Acros organics] and 15 g/L NaCl [Fisher chemical]), 0.1 mM CaCl_2_(H_2_O)_2_ [Acros organics], 1 mM MgCl_2_(H_2_O)_6_ [Fisher bioreagents], 3 µM FeCl_3_ [MP biomedicals], 10 µg/mL thiamine chloride [Acros organics], 25 µg/mL uracil [Acros organics] and 1% glucose [Acros organics]). Overnight cultures (glass tubes, 4 mL LB) were incubated for 16-18h aerobically while shaking (250 RPM) at 30°C (in case of strains containing a *clpB* deletion or a temperature sensitive plasmid) or 37°C. Exponential cultures were made by 1/100 or 1/100 000 (for ABglu) dilution from overnight culture and grown for 3h (for LB) or 28h (for ABglu) at the same shaking speed (250 RPM) and at 37°C or 30°C in 4 mL (LB or ABglu) or 25 mL (Abglu) medium in fitting glass or plastic recipients depending on the strain and the experimental conditions. When appropriate, the following chemicals were added at the indicated final concentration: 50 µg/mL kanamycin sulphate [Panreac Applichem], 100 µg/mL ampicillin sodium salt [Fisher Bioreagents], 30 µg/mL chloramphenicol [Acros organics], 200 ng/mL anhydrotetracycline hydrochlorine [Cayman Chemical Company], 100 µM cuminic acid (4-isopropyl-benzoic acid) [Aldrich chemistry], 2 mM Adenosine 3′,5′-cyclic monophosphate sodium salt (cAMP) [EMD Millipore corp., Merck] and 0.2% L-arabinose [Acros organics].

### Plasmid construction

Plasmids were constructed via Gibson assembly [21], and transformed to the appropriate bacterial strains via electroporation. Primers were ordered at IDT [Leuven, Belgium], and all PCR reactions for construction purposes were performed with Q5 high-fidelty DNA polymerase [New England Biolabs].

The pKD46BB-*syfp2-frt-cat-frt* plasmid was constructed by a two fragment Gibson assembly to couple the *syfp2* gene to a flippable antibiotic marker to provide a selection marker for genomic integration of the *syfp2* gene. The pKD46BB-*frt-cat-frt* plasmid was opened by PCR using primers P1 and P2. The *syfp2* gene was PCR amplified with primers P3 and P4 from strain MG1655 *rph+ lac+ AmalI::frt gtrA::P58-mCherry: parB-SpR codA::parBMt1-1ac0sym-lacOCS-frt recA-recX::recAmwg-syfp2-frt-kan-frt* [22], [23].

To construct the pAJM657-2PA plasmid, the pAJM657 plasmid [which already contains the P*_cym_* promoter and its cognate *cymR^AM^* repressor gene; [24]] was fitted with the genetic parts described below. Parts from the marionette sensor collection [24] include all promotors (except P*_lacI_*), RiboJ insulators, all ribosome binding sites (except the *lacI* RBS), terminators L3S3P21, L3S2P21 and IOT. Terminator ECK120033737 was taken from Chen et al. and synthetically constructed with two overlapping primers [25]. The *tetR* gene and the *rrnB* T1 and T7 terminators are derived from the Tn5-PLtetO-1-*msfGFP* transposon [26], where promotor P*_N25_* and RBSI have been replaced with P*_lacI_* and RBS *lacI* from *E. coli* MG1655 respectively. For the fusion proteins, *T25* and *T18* moieties stem from the *B. pertussis cyaA* gene from plasmids pEB1029 and pEB1030 [27], *mScarlet-I* stems from plasmid pBAM-1-*mScarlet-I* [28], while *mCerulean3* and *cI-78^EP8^*stem from ptrc99A-*mCer-cI-78^EP8^* [16]. These two fluorescent proteins were chosen because of spectral compatibility, distant relatedness and true monomer nature (to avoid weak (inter)dimer interactions) [14], [15]. Finally, linkers ‘DYKDDDDK’ (linking enzymatic (T18/T25) domain and fluorescent domain) and ‘GSGSGS’ (linking fluorescent domain with the cI-78^EP8^ aggregative domain) were derived from Hopp et al. and Chen et al. respectively [29], [30]. Finally, the P*_tet_* promoter was obtained from the pAJM011 plasmid [24]. The fully annotated sequence of the resulting pAJM657-2PA plasmid is provided in Extended Data Fig. 1.

All constructed plasmids were validated by Dreamtaq PCR [Dreamtaq green DNA polymerase kit, Thermo scientific] with dedicated primers (see Extended Data Table 2) and subsequently Sanger-sequenced [Macrogen, Amsterdam, The Netherlands] to verify the entire inserted sequences. Plasmid pAJM657-2PA was fully sequenced with Nanopore whole plasmid sequencing [Macrogen, Amsterdam, The Netherlands].

### Strain construction

The *E. coli* MG1655 *ΔcyaA ΔclpB lacY-syfp2* strain was constructed by three consecutive genomic integrations. First, a transcriptional fusion of the *lacY* gene with *syfp2* was constructed in the MG1655 strain. The *syfp2-frt-cat-frt cassette*, featuring *syfp2* with a flippable chloramphenicol marker, was amplified from the pKD46BB-*syfp2-frt-cat-frt* plasmid with primers P5 and P6. These primers introduce the BBa_B0034 RBS (sequence AAAGAGGAGAA [31]) to ensure independent translation of *syfp2*, and also feature 50 bp complementary overhang to integrate the construct in the *E. coli* genome 5 bp downstream of the *lacY* gene using recombineering as described by Datsenko & Wanner [32]. The *syfp2-frt-cat-frt* cassette was then transformed by electroporation to MG1655 fitted with the pKD46 plasmid that encodes the Lambda red genes needed for homologous recombination leading to genome integration of the cassette [32]. After genomic integration of the *syfp2-frt-cat-frt* cassette and subsequent curing of temperature-sensitive pKD46, the chloramphenicol resistance marker was removed by transiently equipping the MG1655 *lacY-syfp2-frt-cat-frt* strain with pCP20, encoding the Flp recombinase for site-specific recombination at the *frt* sites. The resulting MG1655-*lacY-syfp2-frt* strain was then transformed with pKD46, necessary for construction of *cyaA* and *clpB* gene deletions.

Both *cyaA* and *clpB* gene deletions were consecutively made by recombineering as well [32], [33]. In frame deletions in the *cyaA* and *clpB* genes were constructed by replacing these genes with an *frt-kan-frt* cassette, featuring a flippable kanamycine resistance marker. This *frt-kan-frt* cassette was amplified from plasmid pKD13 [32] using primer pairs P7/P8 for the *cyaA* deletion, and P9/P10 for the *clpB* deletion. The kanamycine resistance marker was removed by Flp recombinase as described earlier. All strains featuring a *cyaA* deletion were at all times complemented with 2 mM cAMP to avoid suppressor mutant emergence. During strain construction, all constructs were verified via Dreamtaq PCR [Dreamtaq green DNA polymerase kit, Thermo scientific] with dedicated primers (see Extended Data Table 2) and subsequently Sanger-sequenced [Macrogen, Amsterdam, The Netherlands].

### Statistical analysis and quantitative data collection

Statistical analysis (Kruskal-Wallis test, and Conover-Iman test for pairwise comparisons between datasets/experimental conditions with Bonferroni correction) was executed with the open-source software R V4.1.1 (R Core Team, 2021) [44]. Starting from a p-value ≤ 0.05, statistical comparisons between cell subpopulations and/or conditions were regarded as significant. Total cell divisions were determined for the PA bearing and PA free daughter cells from the initial PA bearing mother cell during 48 hour long time-lapse fluorescence microscopy experiments by manual monitoring and counting every cell division. A cell division was considered to be complete when two cells could clearly be distinguished from one another in the phase channel. Non-growing cells or cells that could not be properly tracked due to their growth within (rather than around) the microcolony (uncountable) were omitted from all datasets. Further, to obtain the cured T18+T25 dataset, cells containing a PA in both cell poles (double PA) were cured from the complete T18+T25 dataset, as well as cells without a PA (PA-) and cells that stop dividing during the time-lapse due to a too large PA. In fact, too large PAs might cause cell division perturbations and may therefore end up in anucleate cells [34]. In the non-induced (NI) dataset, all cells are PA free and therefore one of the two cell poles was randomly chosen as the PA bearing pole for division counting purposes. The T18, T25, clpB and ΔclpB datasets were cured in the same way as the T18+T25 dataset. Extended Data Fig. 2A and Extended Data Table 3 show uncured datasets visualized in the same way as in Fig. 1C and numbers/percentages of every subpopulation in all of the datasets, respectively.

PA size was measured by manually scoring the combined fluorescent intensity of mScarlet-I and mCerulean3 in the polar focus inside every individual cell. This was measured with the NIS elements Ar analysis 5.20 software (Nikon, Champigny-sur-Marne, France) by drawing a region of interest (ROI), with a surface of 1.01 µm^2^, in the center of the fluorescent focus at the first timepoint of the time-lapse. In case no fluorescent focus was formed, fluorescence was measured in one of the two cell poles (randomly allocated) and for cells with two fluorescent foci in each cell pole, the focus with the highest total fluorescence was scored. Relative PA size was determined by normalizing all PA size values to the highest PA size value per replicate.

All quantitative data originates from three independent experiments. Fig. 1C, Fig. 2C and Extended Data Figs. 2, 4 and 5 were conceived in Graphpad v9.4.1 and v10.0.0 [43].

### Time-lapse fluorescence microscopy

Before microscopic imaging, a 1 mL aliquot from each exponential culture was washed four times with 1 mL 0.85% KCl [Honeywell] (5 min, 3824 RCF) to remove inducers and residual glucose from the permissive ABglu medium. The washed cultures were subsequently diluted to an appropriate OD that allows for long-term single cell monitoring. An aliquot of the diluted culture was added to a cut 2% agarose [Eurogentec, Seraing, Belgium] pad containing the non-permissive ABc medium (ABglu in which 1% glucose was replaced by 1% acid hydrolysed casein (casamino acids) [Neogen]) supplied with 500 µM isopropyl-beta-D-thiogalactoside (IPTG) [Acros organics, dioxiane free]. IPTG ensures proper expression of the *lacY-syfp2* transcriptional fusion when cAMP is present. In fact, the *lac* operon can only be properly expressed when both IPTG and active Cap (activated by cAMP) are available [20]. The ABc IPTG agarose was poured within a 125 µl Geneframe [Thermo Fisher Scientific] which was placed on a microscopy slide [RS France] and covered, after cell loading, with a cover slide [Epredid]. Images taken during time-lapse fluorescence microscopy were captured with a Ti-Eclipse inverted microscope [Nikon, Champigny-sur-Marne, France], equipped with a X60 Plan Apo λ oil objective, a TI-CT-E motorized condenser and a Nikon DS-Qi2 camera. As a light source, the SpecraX LED illuminator [Lumencor, Beaverton, OR, USA] was used and a cage incubator [Okolab, Ottaviano, Italy] enabled stable temperature control during microscopy experiments at 30°C. Fluorescent proteins mScarlet-I, SYFP2 and mCerulean3 were visualized with a triple-edge dichroic (475/540/600 nm). Microscopy images were viewed and acquired with Nikon NIS elements Ar 4.51 and NIS elements Ar analysis 5.20 software [Nikon, Champigny-sur-Marne, France] and subsequently processed with open source software Fiji [35].

### Growth and spot assay

An automated microplate spectrophotometer [Multisckan FC with incubator and SkanIT RE 5.0 software, Thermo Fisher] was used to generate growth curves. Cultures prepared in the same way as for microscopy were diluted to an OD_620_ of 0.05 with an OD meter [Ultrospec 10 cell density meter, Biochrom, Cambourne, UK] and diluted 1/100 in 200 µl permissive (ABglu) or non-permissive (ABc) medium in a 96-well plate [Cellstar, Greiner Bio One] and enclosed with a 120X80 mm transparent plate seal [Greiner]. OD_620_ was measured every 15 minutes in between 30 second shake periods for 72h at 30°C. Three biological replicates and three technical replicates per biological replicate were included and averaged to obtain final growth curves. For every replicate, a 10-times dilution series was spotted (5µl) on solid LB medium to determine starting amount of cells within the microtiter plate wells and incubated at 30°C for 24h.

### Individual based modelling

See Extended Data.

### Image analysis

Fluorescence microscopy images of a representative microcolony were segmented using Ilastik v1.4.0.post1 [36], after which cell tracking was performed using the Strack algorithm [37]. The resulting output was further processed using Python. For each time frame, the amount of progeny generated by each cell in the following 900 minutes was determined. This parameter was then used to generate a color-coded overlay, showcasing the asymmetric inheritance of growth potential. To increase visibility, this progeny-parameter was clipped at a value of 10, as such ensuring the visibility of small growth differences in the presence of fast-growing outliers.

## ACKNOWLEDGEMENTS

This work was funded by doctoral fellowships (11J6222N and 11J6224N to R.V.E.; and 11Q4T24N to K.B.) and project grant (G0D8220N) from the Research Foundation-Flanders (FWO-Vlaanderen).

## AUTHOR CONTRIBUTIONS

R.V.E. and A.A. conceptualized the project and designed experiments. R.V.E. and SJ performed experiments and data analysis. D.O. performed the Individual Based Modeling. R.VE. and K.B. performed image analysis. R.V.E. and A.A. drafted the manuscript and all authors edited the manuscript.

## EXTENDED DATA

**Extended Data Table 1.**
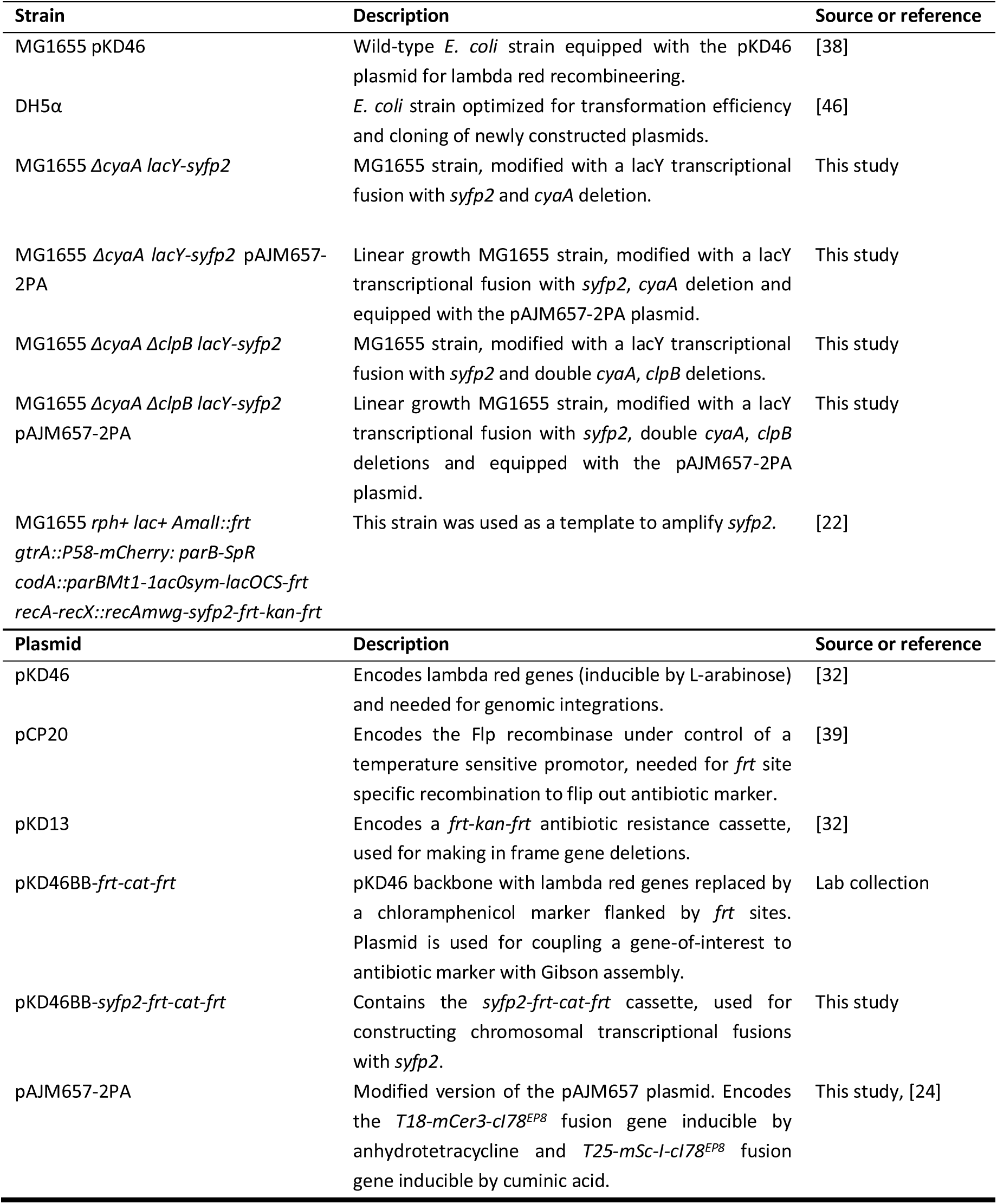
Overview of strains and plasmids used in this study.

| Strain | Description | Source or reference |
| --- | --- | --- |
| MG1655 pKD46 | Wild-type *E. coli* strain equipped with the pKD46 plasmid for lambda red recombineering. | [38] |
| DH5α | *E. coli* strain optimized for transformation efficiency and cloning of newly constructed plasmids. | [46] |
| MG1655 Δ*cyaA* *lacY-syfp2* | MG1655 strain, modified with a *lacY* transcriptional fusion with *syfp2* and *cyaA* deletion. | This study |
| MG1655 Δ*cyaA* *lacY-syfp2* pAJM657-2PA | Linear growth MG1655 strain, modified with a *lacY* transcriptional fusion with *syfp2*, *cyaA* deletion and equipped with the pAJM657-2PA plasmid. | This study |
| MG1655 Δ*cyaA* Δ*clpB* *lacY-syfp2* | MG1655 strain, modified with a *lacY* transcriptional fusion with *syfp2* and double *cyaA*, *clpB* deletions. | This study |
| MG1655 Δ*cyaA* Δ*clpB* *lacY-syfp2* pAJM657-2PA | Linear growth MG1655 strain, modified with a *lacY* transcriptional fusion with *syfp2*, double *cyaA*, *clpB* deletions and equipped with the pAJM657-2PA plasmid. | This study |
| MG1655 *rph+* *lac+* *Amall::frt* *gtrA::P58-mCherry*: *parB-SpR* *codA::parBmt1-1acOsym-lacOCS-frt* *recA-recX::recAmwg-syfp2-frt-kan-frt* | This strain was used as a template to amplify *syfp2*. | [22] |
| Plasmid | Description | Source or reference |
| pKD46 | Encodes lambda red genes (inducible by L-arabinose) and needed for genomic integrations. | [32] |
| pCP20 | Encodes the Flp recombinase under control of a temperature sensitive promotor, needed for *frt* site specific recombination to flip out antibiotic marker. | [39] |
| pKD13 | Encodes a *frt-kan-frt* antibiotic resistance cassette, used for making in frame gene deletions. | [32] |
| pKD46BB-*frt-cat-frt* | pKD46 backbone with lambda red genes replaced by a chloramphenicol marker flanked by *frt* sites. Plasmid is used for coupling a gene-of-interest to antibiotic marker with Gibson assembly. | Lab collection |
| pKD46BB-*syfp2-frt-cat-frt* | Contains the *syfp2-frt-cat-frt* cassette, used for constructing chromosomal transcriptional fusions with *syfp2*. | This study |
| pAJM657-2PA | Modified version of the pAJM657 plasmid. Encodes the *T18-mCer3-cl78<sup>EP8</sup>* fusion gene inducible by anhydrotetracycline and *T25-mSc-I-cl78<sup>EP8</sup>* fusion gene inducible by cuminic acid. | This study, [24] |

**Extended Data Table 2.**
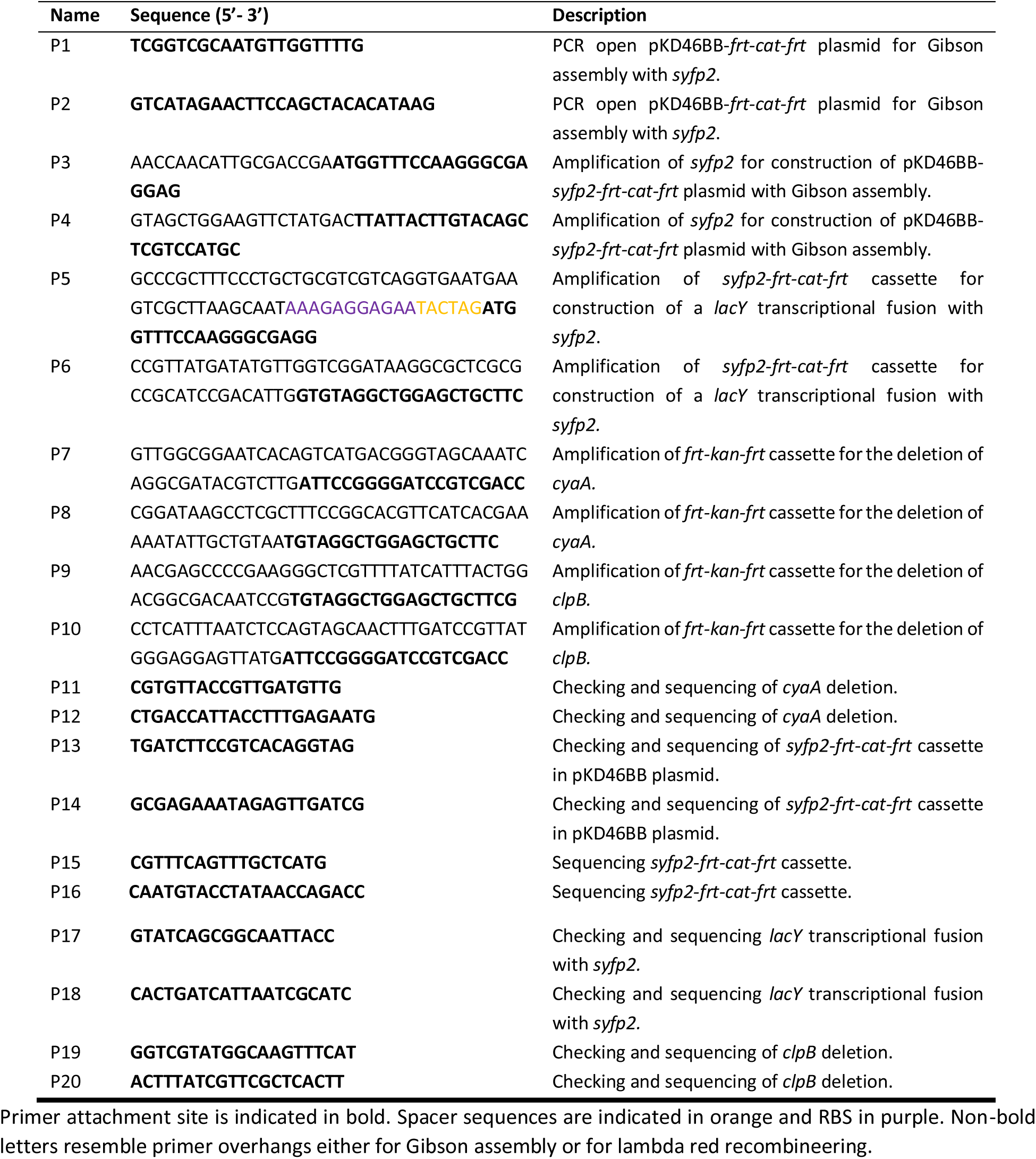
Overview of primers used in this study.

**Extended Data Table 3.** Overview of subpopulation cell numbers (percentages) in every dataset used in this study.

| Dataset | Non-growing | Uncountable | Double PA | PA- | Too large PA | Cured | Total |
| --- | --- | --- | --- | --- | --- | --- | --- |
| **T18+T25** | 9 (3%) | 41 (13%) | 25 (8%) | 19 (6%) | 44 (14%) | 172 (56%) | 310 |
| **NI** | 17 (7%) | 0 (0%) | / | 214 (93%) | / | 214 (93%) | 231 |
| **T18** | 13 (7%) | 0 (0%) | 48 (26%) | 3 (1%) | 0 (0%) | 123 (66%) | 187 |
| **T25** | 7 (4%) | 0 (0%) | 0 (0%) | 143 (76%) | 0 (0%) | 38 (20%) | 188 |
| **clpB** | 7 (4%) | 5 (3%) | 11 (7%) | 17 (11%) | 6 (4%) | 111 (71%) | 157 |
| **ΔclpB** | 7 (4%) | 18 (10%) | 4 (2%) | 13 (8%) | 17 (10%) | 113 (66%) | 172 |

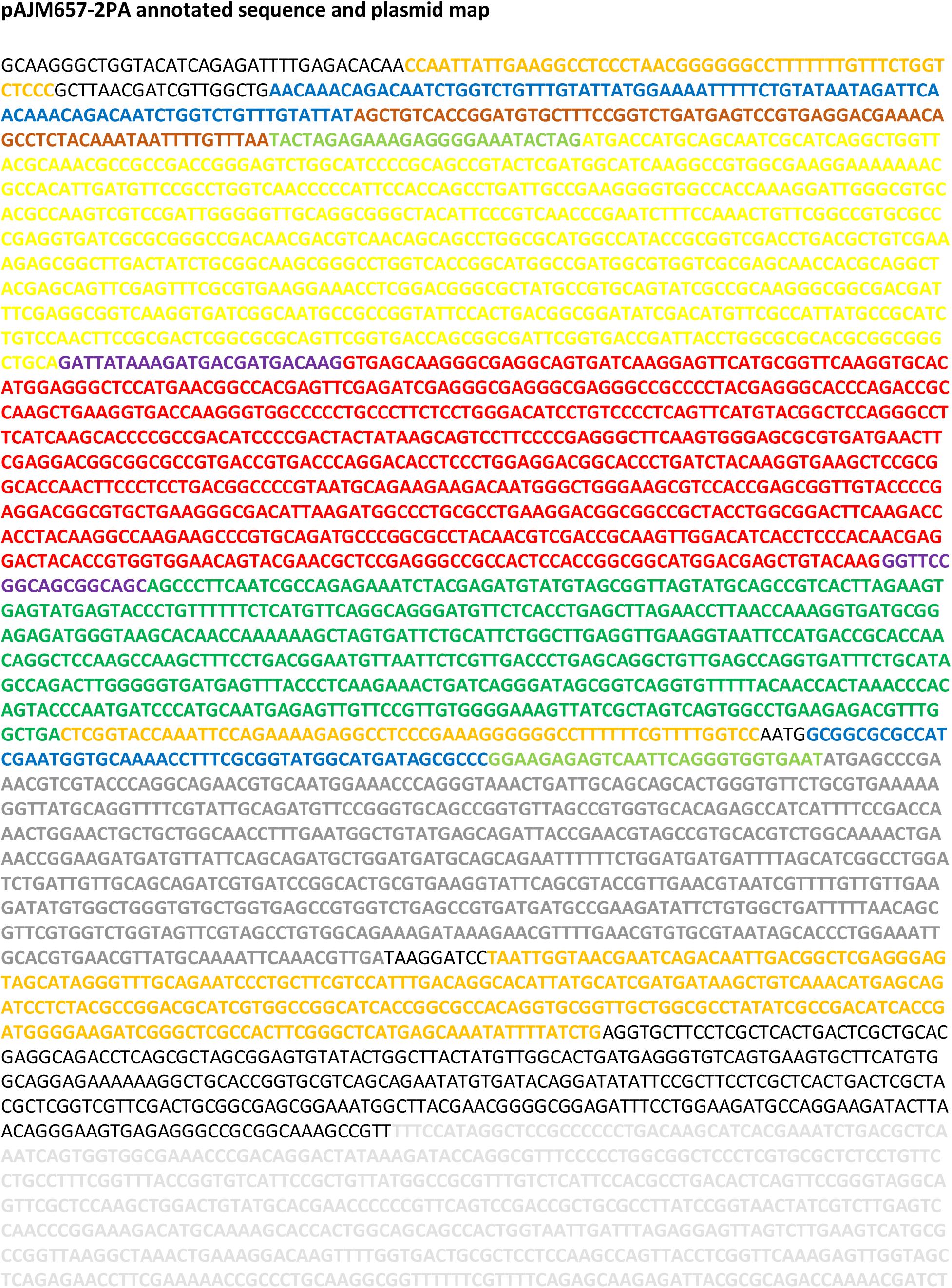
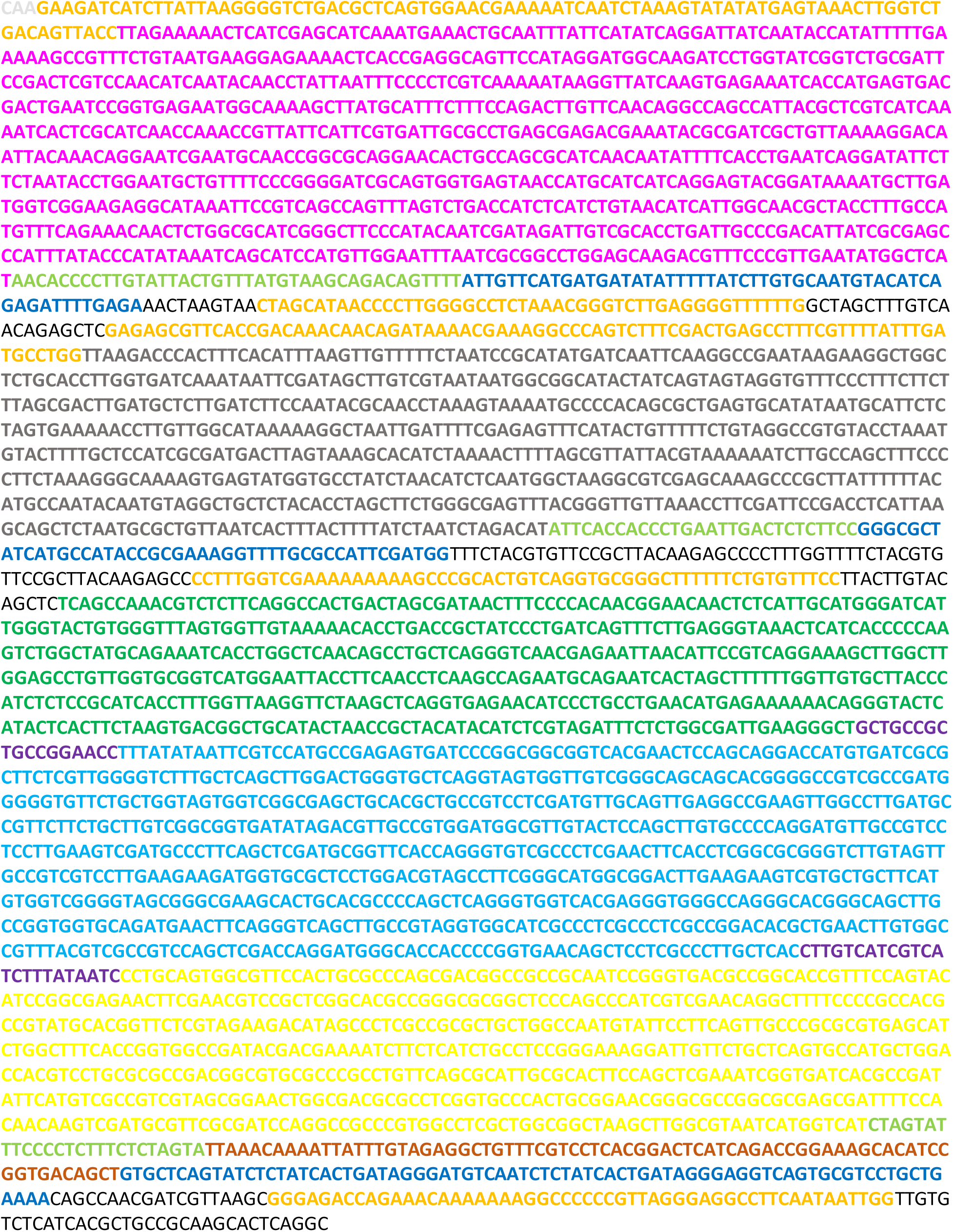

**Extended Data Figure 1:**
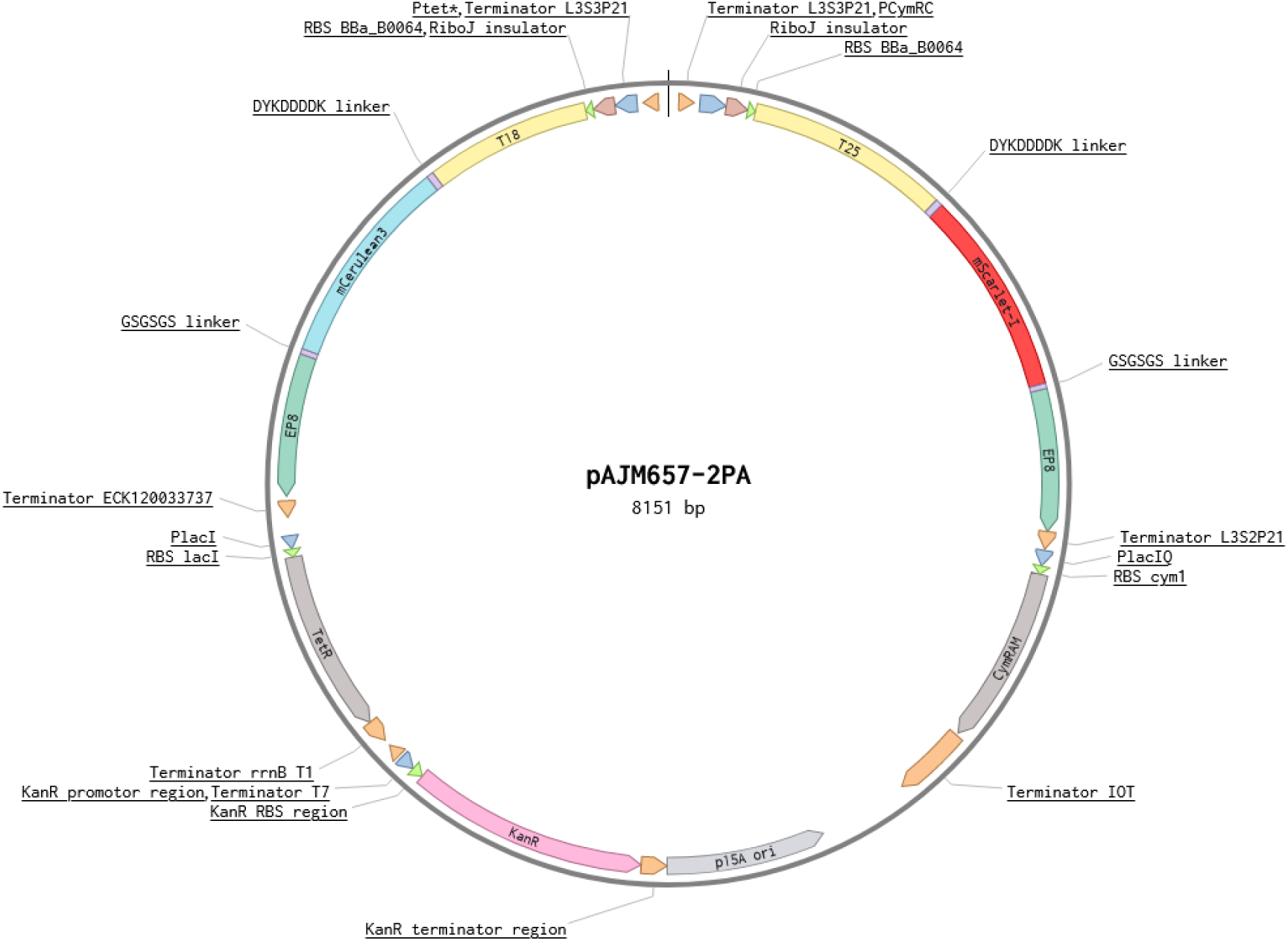
Annotated plasmid map of pAJM657-2PA. Annotations are visualized with following colors: orange for terminators, dark blue for promotors, brown for insulators, light green for ribosome binding sites (RBS), yellow for *T18* and *T25 cyaA* catalytic domain halves, purple for linkers, cyan for *mCerulean3*, red for *mScarlet-I*, green for *cI78^EP8^,* dark grey for repressors, light grey for origin of replication (*p15a ori*), pink for antibiotic marker, and non-bold black for spacers or plasmid backbone sequence. Colors are the same for the annotated sequence and the plasmid map. As the promotor, RBS and terminator of the *kan^R^* gene is unknown, the general region is shown where these genetic parts are predicted. Direction of annotations on plasmid map show direction of genetic parts. Plasmid map was made in Benchling [45].

**Extended Data Figure 2:**
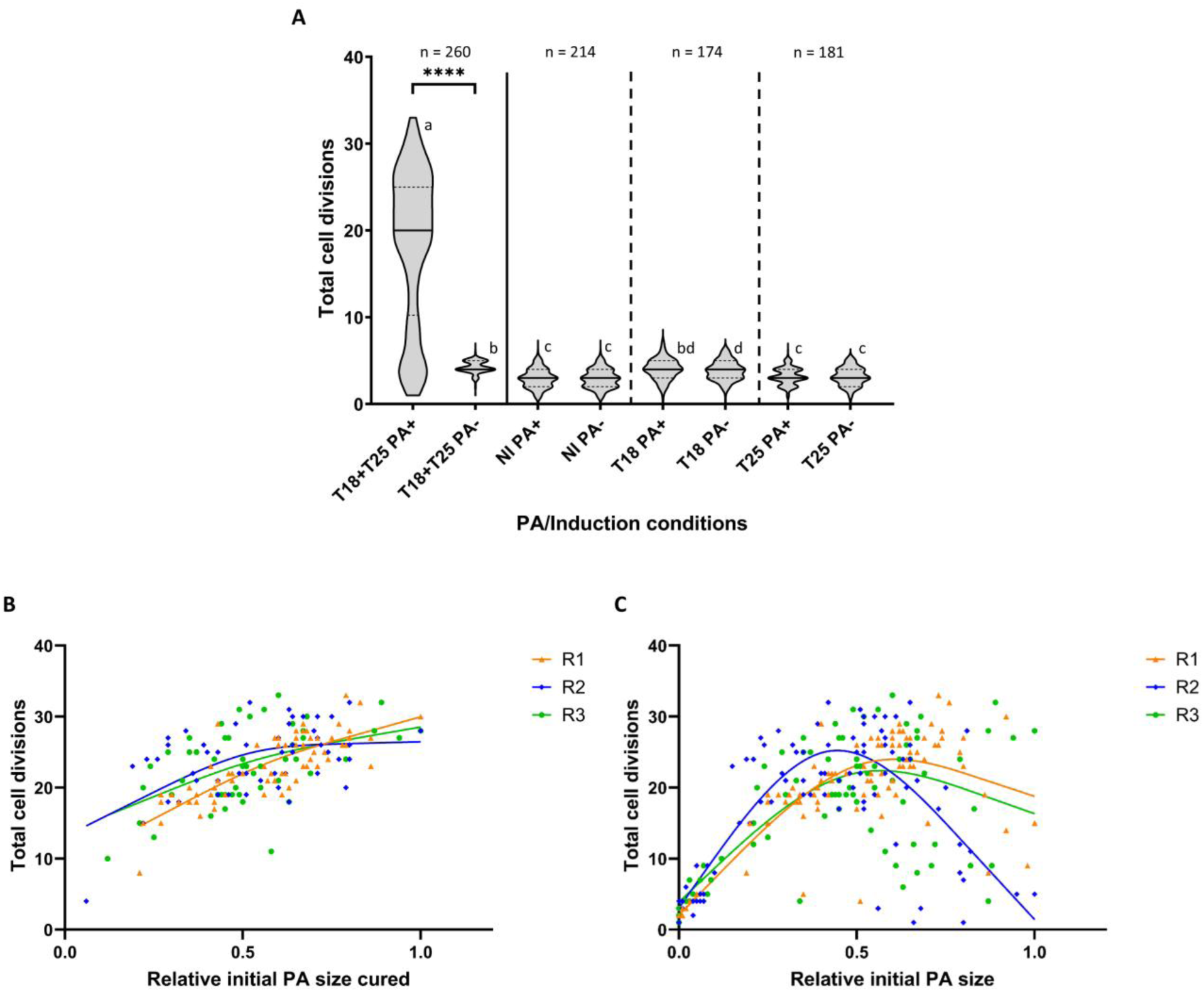
(A) Counted total cell divisions for individual cells in a 48h TLFM experiment on non-permissive (ABc) medium per PA/induction condition displayed as truncated violin plots (uncured datasets). PA+ indicates daughter cells inheriting the PA bearing cell pole whereas PA- indicates daughter cells inheriting the PA free cell pole from the first PA+ mother cell. T18-mCer3-cI78^EP8^ and T25-mSc-I-cI78^EP8^ fusion proteins were either both (T18+T25) or separately (T18/T25) induced or not induced (NI). The number of tracked cells is indicated by n. Violin plots with non-corresponding letters are significantly different from one another (p-value ≤ 0.05, determined by Conover-Iman test with Bonferroni correction after rejection of the null hypothesis of a Kruskal-Wallis test). Four asterisks indicate a p-value ≤ 0.0001. (B and C) Scatterplot correlating the total number of divisions of a cell with its initial PA size (normalized to the largest PA size in the dataset per replicate, at the first timepoint of a 48h time-lapse) on non-permissive (ABc) medium for MG1655 *ΔcyaA ΔclpB lacY-syfp2* pAJM657-2PA with pulse-induction of the T18-mCer3-cI78^EP8^ and T25-mSc-I-cI78^EP8^ fusion proteins. Replicates indicated with orange triangles for R1, blue diamonds for R2 and green dots for R3 and a trendline for each replicate in the same color (spline with 3 knots). Both cured (B, n = 172) and uncured (C, n = 260) datasets are shown for comparison.

**Extended Data Figure 3:**
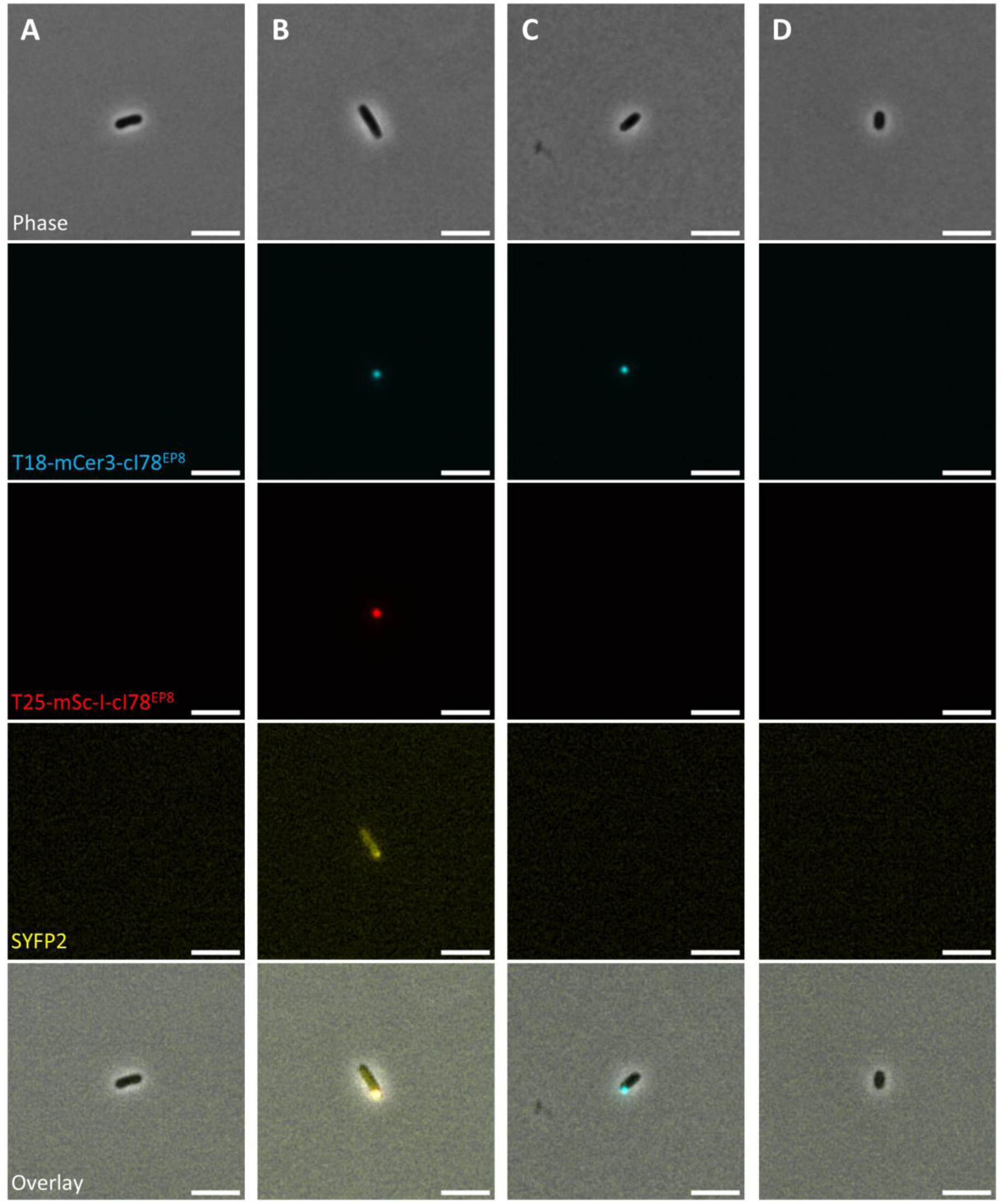
Representative fluorescence microscopy images of representative MG1655 *ΔcyaA ΔclpB lacY-syfp2* pAJM657-2PA cells 1,5h after either (A) no induction, (B) pulse induction of both T18-mCer3-cI78^EP8^ and T25-mSc-I-cI78^EP8^, (C) pulse induction of only T18-mCer3-cI78^EP8^, and (D) pulse induction of only T25-mSc-I-cI78^EP8^. Scale bar corresponds to 5 µm.

**Extended Data Figure 4:**
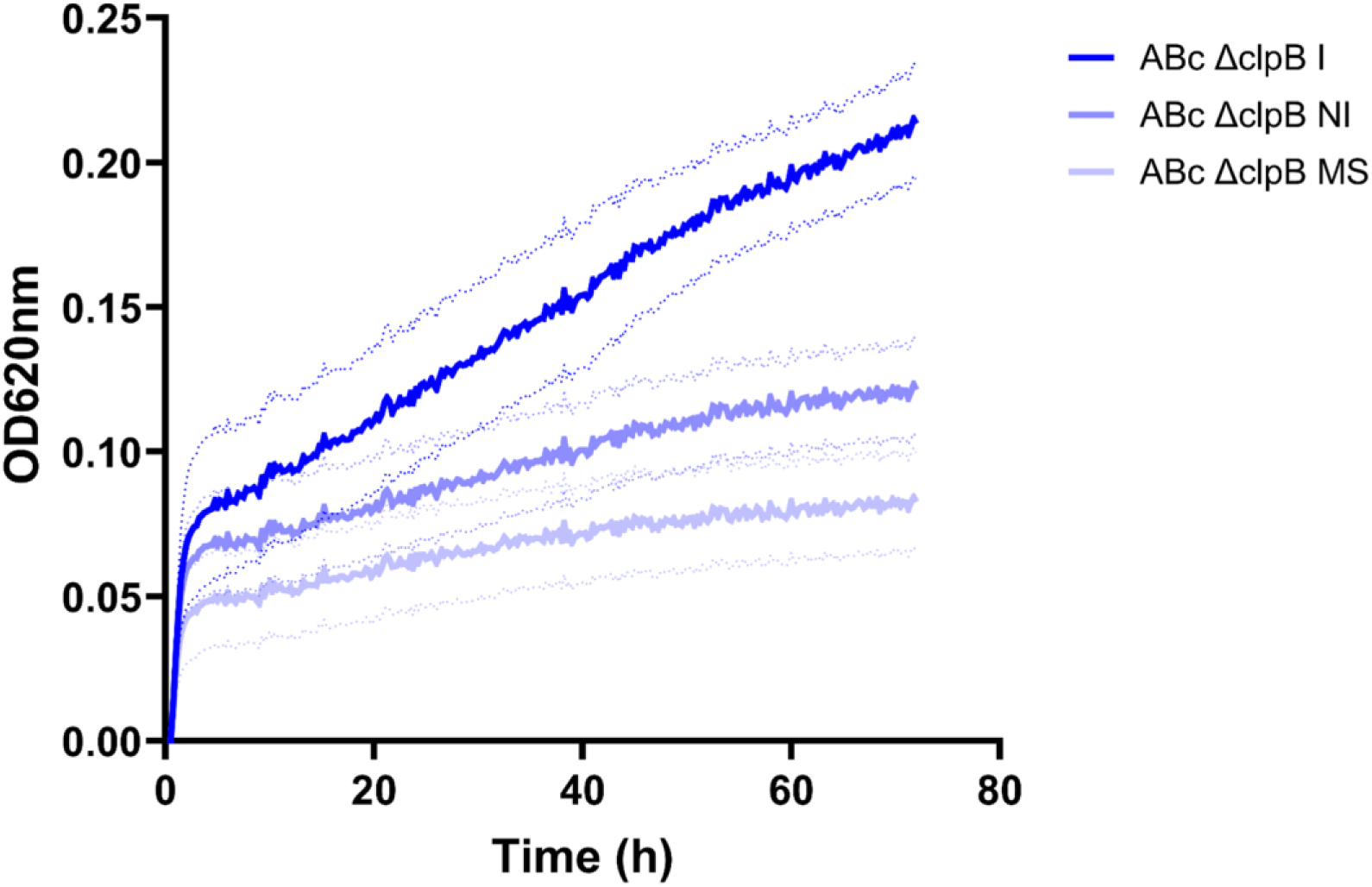
Optical density (OD_620_) monitored bulk growth profile of MG1655 *ΔcyaA ΔclpB lacY-syfp2* pAJM657-2PA cells either induced with prior pulse-induction of both T18-mCer3-cI78^EP8^ and T25-mSc-I-cI78^EP8^ (I) or non-induced (NI), or MG1655 *ΔcyaA ΔclpB lacY-syfp2* cells (lacking the pAJM657-2PA plasmid; MS) in non-permissive (ABc) medium, together with the corresponding standard deviation (dotted lines).

**Extended Data Figure 5:**
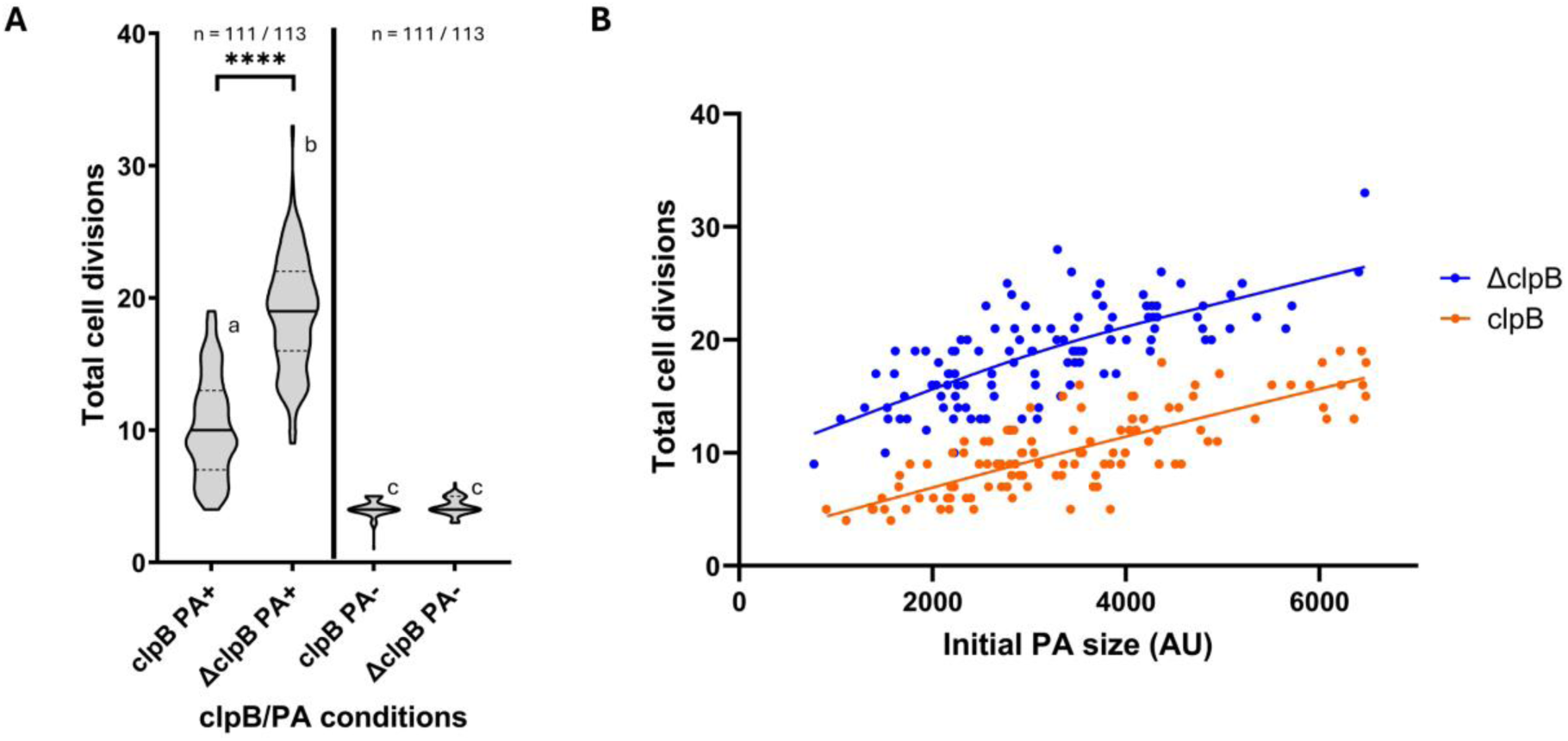
(A) Counted total cell divisions for individual cells in a 48h TLFM experiment on non-permissive (ABc) medium per clpB/PA condition displayed as truncated violin plots (cured datasets). PA+ indicates daughter cells inheriting the PA bearing cell pole, whereas PA- indicates daughter cells inheriting the PA free cell pole from the first PA+ mother cell. Both T18-mCer3-cI78^EP8^ and T25-mSc-I-cI78^EP8^ fusion proteins were either pulse-induced in MG1655 *ΔcyaA ΔclpB lacY-syfp2* pAJM657-2PA (ΔclpB) or MG1655 *ΔcyaA lacY-syfp2* pAJM657-2PA (clpB). The number of tracked cells is indicated by n. Violin plots with non-corresponding letters are significantly different from one another (p-value ≤ 0.05, determined by Conover-Iman test with Bonferroni correction after rejection of the null hypothesis of a Kruskal-Wallis test). Four asterisks indicate a p-value ≤ 0.0001. (B) Scatterplot correlating the total number of divisions of a cell with its initial PA size (in arbitrary units, at the first timepoint of a 48h time-lapse) on non-permissive (ABc) medium for MG1655 *ΔcyaA ΔclpB lacY-syfp2* pAJM657-2PA (ΔclpB, blue dots and trendline (spline with 3 knots), n = 113) and MG1655 *ΔcyaA lacY-syfp2* pAJM657-2PA (clpB, orange dots and trendline (spline with 3 knots), n = 111) with pulse expressed T18-mCer3-cI78^EP8^ and T25-mSc-I-cI78^EP8^ fusion proteins.

### Individual based modelling

To further examine the theorical basis underlying the linear growth dynamics of the engineered chassis, an individual based model was constructed in which each cell is simulated as a separate entity with its own state variables. The tracked state variables per cell are the cell volume, the amount of an essential protein and the amount of an essential metabolite produced by this protein.

The state variable volume (𝑽) is used to simulate cellular growth. Its value is dimensionless, as it represents the relative volume with respect to the volume of a newly divided cell. When the volume of a cell reaches double that of the initial value, the cell will divide, producing two daughter cells. Each daughter cell will have half the volume of the original cell. Similarly, the amount of essential metabolite will be divided equally among daughter cells. The symmetry of division of the essential protein, however, depends on the parameter 𝒅_𝒔_, which has a value between 0 and 1, and where 0 (or 1) represents full asymmetric division while 0.5 represents full symmetric division. Cells are assumed to have unlimited access to nutrients, thus disregarding growth restrictions related to the media composition. Growth can be restricted, however, based on the amount of essential metabolite. We opted to use the Monod equation [40] to represent the dependency between the growth rate (𝝁) and the concentration of essential metabolite (𝑴). Additionally, the growth rate is equal to zero when the essential metabolite concentration drops below a certain threshold (𝑴_𝑻_). The rate of change of the cellular volume (𝑽) is given in Equation 1. Note that we use the concentration of essential metabolite here, which we can obtain by dividing the amount of essential metabolite by the relative volume of the cell.

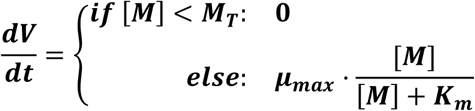

Each cell at the start of the simulation will contain a starting amount of essential protein (𝑬). The production of essential protein is assumed to be constitutive in wild-type cells with a constant rate (𝜶_𝑬_). During linear growth, essential protein is not produced, which is reflected in the simulation by setting the production rate to zero. Essential protein is also assumed to degrade at a constant rate (𝜸_𝑬_). The rate of change of the essential protein (𝑬) is given in Equation 2.

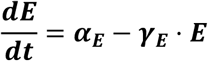

Production of essential metabolite is dependent on the amount of essential protein within the cell and is assumed to be produced at a constant rate (𝜶_𝑬_). Additionally, it is assumed to degrade at a constant rate (𝜸_𝑴_). The rate of change of the essential metabolite (𝑴) is given in Equation 3.

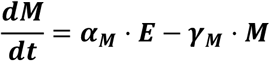

The model is written using the programming language Julia v1.10 [41]), and is simulated as a system of ordinary differential equations (ODEs) using the DifferentialEquations.jl package [42]. The code is freely accessible on github: https://github.com/DriesOome/LinearGrowthIBM.git. As the model is individual based, the size of the system of ODEs increases each time a new cell is born. The solver utilizes a callback system to simulate the discrete division events. Namely, the solver pauses when a cell reaches the division threshold, adds the required equations to the ODE system and modifies the involved state variables to their correct values before restarting the simulation.

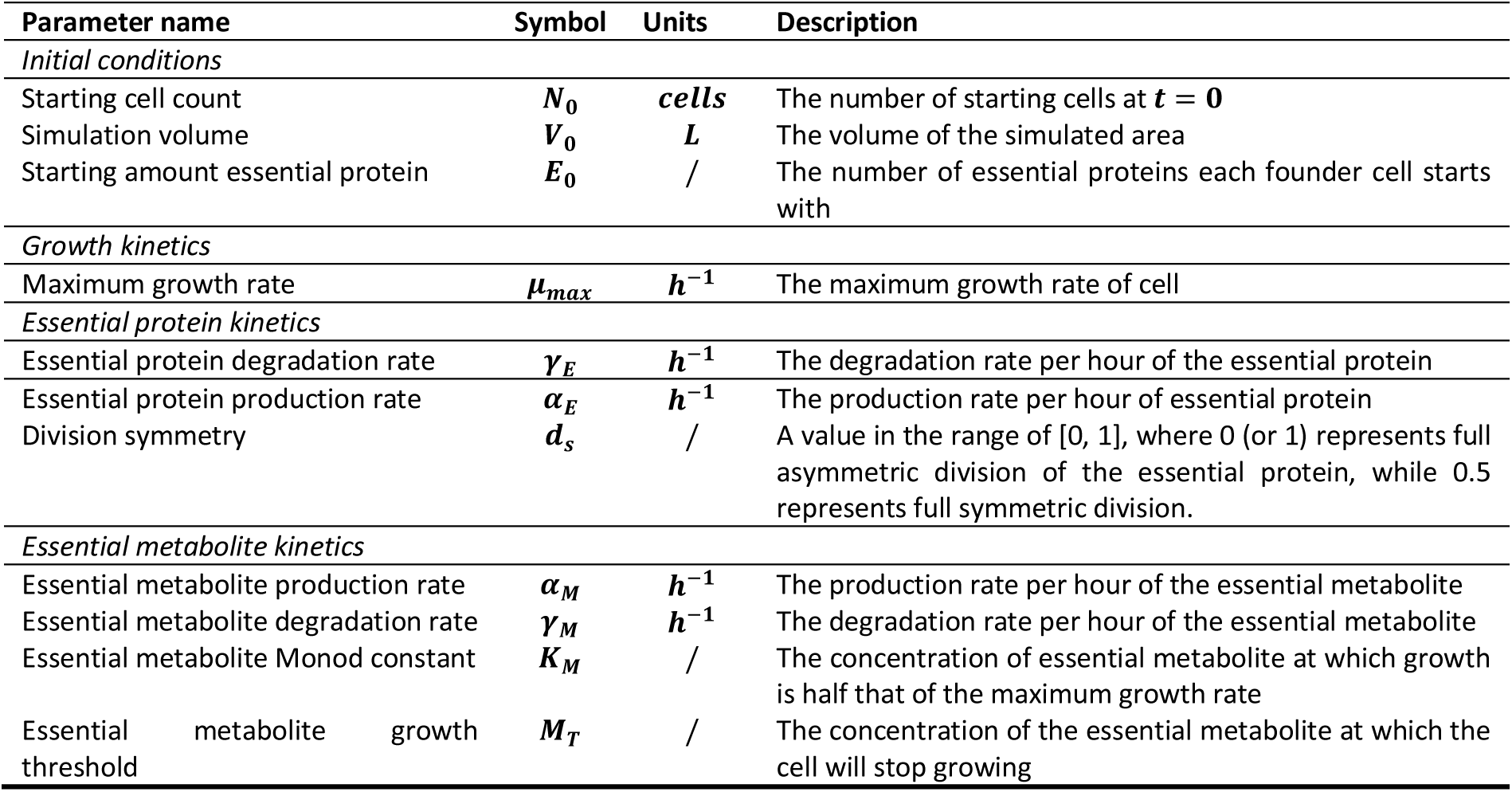

| Parameter name | Symbol | Units | Description |
| --- | --- | --- | --- |
| *Initial conditions* |  |  |  |
| Starting cell count | $N_0$ | *cells* | The number of starting cells at $t = 0$ |
| Simulation volume | $V_0$ | *L* | The volume of the simulated area |
| Starting amount essential protein | $E_0$ | / | The number of essential proteins each founder cell starts with |
| *Growth kinetics* |  |  |  |
| Maximum growth rate | $\mu_{max}$ | $h^{-1}$ | The maximum growth rate of cell |
| *Essential protein kinetics* |  |  |  |
| Essential protein degradation rate | $\gamma_E$ | $h^{-1}$ | The degradation rate per hour of the essential protein |
| Essential protein production rate | $\alpha_E$ | $h^{-1}$ | The production rate per hour of essential protein |
| Division symmetry | $d_s$ | / | A value in the range of [0, 1], where 0 (or 1) represents full asymmetric division of the essential protein, while 0.5 represents full symmetric division. |
| *Essential metabolite kinetics* |  |  |  |
| Essential metabolite production rate | $\alpha_M$ | $h^{-1}$ | The production rate per hour of the essential metabolite |
| Essential metabolite degradation rate | $\gamma_M$ | $h^{-1}$ | The degradation rate per hour of the essential metabolite |
| Essential metabolite Monod constant | $K_M$ | / | The concentration of essential metabolite at which growth is half that of the maximum growth rate |
| Essential metabolite growth threshold | $M_T$ | / | The concentration of the essential metabolite at which the cell will stop growing |

## REFERENCES

[1] T. Nyström, “Stationary-phase physiology,” Annu. Rev. Microbiol., vol. 58, pp. 161–181, 2004.

[2] A. Meunier, F. Cornet, and M. Campos, “Bacterial cell proliferation: From molecules to cells,” FEMS Microbiol. Rev., vol. 45, no. 1, pp. 1–21, 2021.

[3] J. M. Gonzalez and B. Aranda, “Microbial Growth under Limiting Conditions-Future Perspectives,” Microorganisms, vol. 11, no. 7, pp. 1–21, 2023.

[4] M. M. Logsdon and B. B. Aldridge, “Stable regulation of cell cycle events in mycobacteria: Insights from inherently heterogeneous bacterial populations,” Front. Microbiol., vol. 9, no. MAR, pp. 1–15, 2018.

[5] B. B. Aldridge et al., “Asymmetry and Aging of Mycobacterial Cells Lead to Variable Growth and Antibiotic Susceptibility,” Science, vol. 529, no. 2006, pp. 100–104, 2011.

[6] D. Oh, Y. Yu, H. Lee, J. H. Jeon, B. L. Wanner, and K. Ritchie, “Asymmetric polar localization dynamics of the serine chemoreceptor protein Tsr in Escherichia coli,” PLoS One, vol. 13, no. 5, pp. 1–12, 2018.

[7] G. Ebersbach, A. Briegel, G. J. Jensen, and C. Jacobs-Wagner, “A Self-Associating Protein Critical for Chromosome Attachment, Division, and Polar Organization in Caulobacter,” Cell, vol. 134, no. 6, pp. 956–968, 2008.

[8] A. B. Lindner, R. Madden, A. Demarez, E. J. Stewart, and F. Taddei, “Asymmetric segregation of protein aggregates is associated with cellular aging and rejuvenation,” Proc. Natl. Acad. Sci. U. S. A., vol. 105, no. 8, pp. 3076–3081, 2008.

[9] N. V. Mushnikov, A. Fomicheva, M. Gomelsky, and G. R. Bowman, “Inducible asymmetric cell division and cell differentiation in a bacterium,” Nat. Chem. Biol., vol. 15, no. 9, pp. 925–931, 2019.

[10] B. Görke and J. Stülke, “Carbon catabolite repression in bacteria: Many ways to make the most out of nutrients,” Nat. Rev. Microbiol., vol. 6, no. 8, pp. 613–624, 2008.

[11] M. H. Saier, “Multiple Mechanisms Controlling Carbon Metabolism in Bacteria,” Biotechnology, vol. 235, no. 6, pp. 1–5, 1998.

[12] G. Karimova, J. Pidoux, A. Ullmann, and D. Ladant, “A bacterial two-hybrid system based on a reconstituted signal transduction pathway,” Proc. Natl. Acad. Sci. U. S. A., vol. 95, no. 10, pp. 5752–5756, 1998.

[13] A. Battesti and E. Bouveret, “The bacterial two-hybrid system based on adenylate cyclase reconstitution in Escherichia coli,” Methods, vol. 58, no. 4, pp. 325–334, 2012.

[14] D. S. Bindels et al., “MScarlet: A bright monomeric red fluorescent protein for cellular imaging,” Nat. Methods, vol. 14, no. 1, pp. 53–56, 2016.

[15] M. L. Markwardt et al., “An improved cerulean fluorescent protein with enhanced brightness and reduced reversible photoswitching,” PLoS One, vol. 6, no. 3, 2011.

[16] S. K. Govers, J. Mortier, A. Adam, and A. Aertsen, “Protein aggregates encode epigenetic memory of stressful encounters in individual escherichia coli cells,” Plos Biol., vol. 16, no. 8. 2018.

[17] S. Lee, M. E. Sowa, J. M. Choi, and F. T. F. Tsai, “The ClpB/Hsp104 molecular chaperone - A protein disaggregating machine,” J. Struct. Biol., vol. 146, no. 1–2, pp. 99–105, 2004.

[18] S. M. Doyle and S. Wickner, “Hsp104 and ClpB: protein disaggregating machines,” Trends Biochem. Sci., vol. 34, no. 1, pp. 40–48, 2009.

[19] P. Katikaridis, V. Bohl, and A. Mogk, “Resisting the Heat: Bacterial Disaggregases Rescue Cells From Devastating Protein Aggregation,” Front. Mol. Biosci., vol. 8, no. May, pp. 1–14, 2021.

[20] M. Lewis, “Allostery and the lac operon,” J. Mol. Biol., vol. 425, no. 13, pp. 2309–2316, 2013.

[21] D. G. Gibson, L. Young, R. Y. Chuang, J. C. Venter, C. A. Hutchison, and H. O. Smith, “Enzymatic assembly of DNA molecules up to several hundred kilobases,” Nat. Methods, vol. 6, no. 5, pp. 343–345, 2009.

[22] J. Wiktor et al., “RecA finds homologous DNA by reduced dimensionality search,” Nature, vol. 597, no. 7876, pp. 426–429, 2021.

[23] G. J. Kremers, J. Goedhart, E. B. Van Munster, and T. W. J. Gadella, “Cyan and yellow super fluorescent proteins with improved brightness, protein folding, and FRET förster radius,” Biochemistry, vol. 45, no. 21, pp. 6570–6580, 2006.

[24] A. J. Meyer, T. H. Segall-Shapiro, E. Glassey, J. Zhang, and C. A. Voigt, “Escherichia coli ‘Marionette’ strains with 12 highly optimized small-molecule sensors,” Nat. Chem. Biol., vol. 15, no. 2, pp. 196–204, 2019.

[25] Y. J. Chen et al., “Characterization of 582 natural and synthetic terminators and quantification of their design constraints,” Nat. Methods, vol. 10, no. 7, pp. 659–664, 2013.

[26] I. Passaris, W. M. Tadesse, E. Gayán, and A. Aertsen, “Construction and validation of the Tn5-PLtetO-1-msfGFP transposon as a tool to probe protein expression and localization,” J. Microbiol. Methods, vol. 161, no. April, pp. 56–62, 2019.

[27] A. Battesti and E. Bouveret, “Improvement of bacterial two-hybrid vectors for detection of fusion proteins and transfer to pBAD-tandem affinity purification, calmodulin binding peptide, or 6-histidine tag vectors,” Proteomics 2008, 8, pp. 4768–4771, 2008.

[28] J. Mortier et al., “Gene erosion can lead to gain-of-function alleles that contribute to bacterial fitness,” MBio, vol. 12, no. 4, pp. 1–21, 2021.

[29] T. P. Hopp et al., “A short polypeptide marker sequence useful for recombinant protein identification and purification,” Bio/Technology, vol. 6, no. 10, pp. 1204–1210, 1988.

[30] X. Chen, J. L. Zaro, and W. C. Shen, “Fusion protein linkers: Property, design and functionality,” Adv. Drug Deliv. Rev., vol. 65, no. 10, pp. 1357–1369, 2013.

[31] Michael B. Elowitz and Stanislas Leibler, “A synthetic oscillatory network of transcriptional regulators,” Nature, vol. 403, pp. 335–338, 2000.

[32] K. A. Datsenko and B. L. Wanner, “One-step inactivation of chromosomal genes in Escherichia coli K-12 using PCR products,” Proc. Natl. Acad. Sci. U. S. A., vol. 97, no. 12, pp. 6640–6645, 2000.

[33] T. Baba et al., “Construction of Escherichia coli K-12 in-frame, single-gene knockout mutants: The Keio collection,” Mol. Syst. Biol., vol. 2, 2006.

[34] J. Mortier et al., “Protein aggregates act as a deterministic disruptor during bacterial cell size homeostasis,” Cell. Mol. Life Sci., vol. 80, no. 12, pp. 1–13, 2023.

[35] J. Schindelin et al., “Fiji: An open-source platform for biological-image analysis,” Nat. Methods, vol. 9, no. 7, pp. 676–682, 2012.

[36] S. Berg et al., “Ilastik: Interactive Machine Learning for (Bio)Image Analysis,” Nat. Methods, vol. 16, no. 12, pp. 1226–1232, 2019.

[37] H. Todorov, T. Miguel Trabajo, and J. R. van der Meer, “STrack: A Tool to Simply Track Bacterial Cells in Microscopy Time-Lapse Images,” mSphere, vol. 8, no. 2, 2023.

[38] F. R. Blattner et al., “The complete genome sequence of Escherichia coli K-12,” Science, vol. 277, no. 5331, pp. 1453–1462, 1997.

[39] P. P. Cherepanov and W. Wackernagel, “Gene disruption in Escherichia coli: TcR and KmR cassettes with the option of Flp-catalyzed excision of the antibiotic-resistance determinant,” Gene, vol. 158, no. 1, pp. 9–14, 1995.

[40] J. Monod, “THE GROWTH OF BACTERIAL CULTURES” *Annu*. Rev. M, vol. 3, no. Xl, pp. 371–394, 1949.

[41] J. Bezanson, A. Edelman, S. Karpinski, and V. B. Shah, “Julia: A fresh approach to numerical computing,” SIAM Rev., vol. 59, no. 1, pp. 65–98, 2017.

[42] C. Rackauckas and Q. Nie, “DifferentialEquations.jl – A Performant and Feature-Rich Ecosystem for Solving Differential Equations in Julia,” J. Open Res. Softw., vol. 5, no. 1, p. 15, 2017.

[43] Truncated violin plots, scatter plots and bulk growth curves were made using GraphPad Prism version 9.4.1 and 10.0.0 for Windows (64-bit), GraphPad Software, Boston, Massachusetts USA, www.graphpad.com

[44] RStudio Team (2021). RStudio: Integrated Development Environment for R. RStudio, PBC, Boston, MA URL http://www.rstudio.com/.

[45] Benchling [Biology Software]. (2024). Retrieved from https://benchling.com

[46] Bethesda Research Laboratories. 1986. BRL pUC host: E. coli DH5α competent cells. Focus 8(2):9

